# Methylene Blue Inhibits In Vitro the SARS-CoV-2 Spike – ACE2 Protein-Protein Interaction – A Mechanism That Can Contribute to Its Antiviral Activity Against COVID-19

**DOI:** 10.1101/2020.08.29.273441

**Authors:** Damir Bojadzic, Oscar Alcazar, Peter Buchwald

**Affiliations:** Diabetes Research Institute, University of Miami, Miami, Florida, USA; Department of Molecular and Cellular Pharmacology, Miller School of Medicine, University of Miami, Miami, Florida, USA

**Keywords:** ACE2, antiviral, chloroquine, COVID-19, methylene blue, protein-protein interaction, SARS-CoV-2, spike protein

## Abstract

Due to our interest in the chemical space of organic dyes to identify potential small-molecule inhibitors (SMIs) for protein-protein interactions (PPIs), we initiated a screen of such compounds to assess their inhibitory activity against the interaction between SARS-CoV-2 spike protein and its cognate receptor ACE2, which is the first critical step initiating the viral attachment and entry of this coronavirus responsible for the ongoing COVID-19 pandemic. As part of this, we found that methylene blue, a tricyclic phenothiazine compound approved by the FDA for the treatment of methemoglobinemia and used for other medical applications (including the inactivation of viruses in blood products prior to transfusion when activated by light), inhibits this interaction. We confirmed that it does so in a concentration-dependent manner with a low micromolar half-maximal inhibitory concentration (IC_50_ = 3 μM) in our protein-based ELISA-type setup, while chloroquine, siramesine, and suramin showed no inhibitory activity in this assay. Erythrosine B, which we have shown before to be a promiscuous SMI of PPIs, also inhibited this interaction with an activity similar, possibly slightly higher, than those found for it for other PPIs. This PPI inhibitory activity of methylene blue could contribute to its antiviral activity against SARS-CoV-2 even in the absence of light by blocking its attachment to ACE2-expressing cells and making this inexpensive and widely available drug potentially useful in the prevention and treatment of COVID-19 as an oral or inhaled medication.

## INTRODUCTION

Severe acute respiratory syndrome-coronavirus 2 (SARS-CoV-2), a novel betacoronavirus and the most recent one of the seven coronaviruses (CoVs) known to infect humans, is responsible for COVID-19, which has been declared a pandemic by the World Health Organization in March 2020 and continues to spread worldwide (Liu et al., 2020; Matheson and Lehner, 2020; Moore and June, 2020). While four CoVs (HCoV 229E, OC43, NL63, and HKU1) are responsible for about one third of the common cold cases in humans, three have caused recent epidemics associated with considerable mortality: SARS-CoV-1 (2002– 2003, causing ∼10% mortality), MERS-CoV (Middle East respiratory syndrome coronavirus; 2012, causing ∼35% mortality), and now SARS-CoV-2 (2019-2020), which seems to be less lethal but more transmissible (Guy et al., 2020). COVID-19 is the most infectious agent in a century (Tiwari et al., 2020) and has already caused infections in the order of tens of millions and deaths that are likely to be in the order of millions worldwide. According to current early estimates, about 3% of infected individuals need hospitalization and 0.5% die, a range that is strongly age-dependent increasing from 0.001% in <20 years old to 8.3% in those >80 years old (Salje et al., 2020). Accordingly, there is considerable interest in possible preventive or therapeutic treatments. There are several possible targets in the coronavirus life cycle for therapeutic interventions including attachment and entry, uncoating, gRNA replication, translation in endoplasmic reticulum (ER) and Golgi, assembly, and virion release (Guy et al., 2020). Viral attachment and entry are particularly promising as they are the first steps in the replication cycle and take place at a relatively accessible extracellular site; hence, they have been explored for intervention purposes for several viruses (Melby and Westby, 2009). CoVs use their glycosylated spike (S) protein to bind to their cognate cell surface receptors and initiate membrane fusion and virus entry. For both SARS-CoV and SARS-CoV-2, the S protein mediates entry into cells by binding to angiotensin converting enzyme 2 (ACE2) via its receptor-binding domain (RBD) followed by proteolytic activation by human proteases (Lan et al., 2020; Matheson and Lehner, 2020; Shang et al., 2020; Sivaraman et al., 2020). Blockade of this RBD–ACE2 protein-protein interaction (PPI) can disrupt infection efficiency; for example, SARS-CoV-2 RBD protein was shown to block S protein mediated SARS-CoV-2 pseudovirus entry into their respective ACE2 receptor-expressing target cells (Tai et al., 2020). Antibodies can be quite effective PPI inhibitors, and they are highly target-specific and relatively stable *in vivo*. However, they cannot reach intracellular targets and, as all other protein therapies, are hindered by problems such as low solubility, propensity for immunogenicity, long elimination half-lives, lack of oral bioavailability, product heterogeneity, and possible manufacturing and storage stability issues. Since they are foreign proteins, they elicit strong immune response in certain patients (Suntharalingam et al., 2006; Wadman, 2006; Leader et al., 2008), and even if approved for clinical use, they tend to have more post-market safety issues than small-molecule drugs (Downing et al., 2017). Small-molecule inhibitors (SMIs) are more challenging to identify for PPIs, but it is now well established that they can be effective against certain PPIs and can offer useful alternatives. There are now >40 PPIs targeted by SMIs that are in preclinical development, and two such SMIs are approved for clinical use (venetoclax and lifitegrast) (Arkin and Wells, 2004; Milroy et al., 2014; Scott et al., 2016; Bojadzic and Buchwald, 2018).

Due to our interest in the chemical space of organic dyes to identify potential SMIs for PPIs (Margolles-Clark et al., 2009a; Margolles-Clark et al., 2009b; Ganesan et al., 2011; Song et al., 2014; Chen et al., 2017; Bojadzic and Buchwald, 2018; Bojadzic et al., 2018), we initiated a screen of such compounds for their ability to inhibit the interaction between SARS-CoV-2 spike protein and its cognate receptor ACE2, which is the first critical step initiating the viral attachment and entry of this CoV. As part of this, we found that methylene blue, a tricyclic phenothiazine compound approved for the treatment of acquired methemoglobinemia and some other uses (Clifton and Leikin, 2003; Schirmer et al., 2011; Bistas and Sanghavi, 2020), inhibits this interaction, and we have confirmed that it does so in a concentration-dependent manner. This can contribute to the antiviral activity of this inexpensive and widely available dye-based drug against SARS-CoV-2 making it potentially useful in the prevention and treatment of COVID-19, especially in non-industrialized nations.

## MATERIALS AND METHODS

### Binding Assays

Methylene blue and all other test compounds used here were obtained from Sigma-Aldrich (St. Louis, MO, USA). ACE2-Fc and SARS-CoV-2 S1 or RBD with His tag proteins used in the binding assay were obtained from SinoBiological (Wayne, PA, USA); catalog no. 10108-H05H, 40592-V08H, and 40591-V08H). Binding inhibition assays were performed in a 96-well cell-free format similar to the one described before (Margolles-Clark et al., 2009b; Ganesan et al., 2011; Song et al., 2014; Chen et al., 2017). Briefly, microtiter plates (Nunc F Maxisorp, 96-well; Thermo Fisher Scientific, Waltham, MA, USA) were coated overnight at 4 °C with 100 μL/well of Fc-conjugated ACE2 receptor diluted in PBS pH 7.2. This was followed by blocking with 200 μL/well of SuperBlock (PBS) (Thermo Fisher Scientific) for 1 h at RT. Then, plates were washed twice using washing solution (PBS pH 7.4, 0.05% Tween-20) and tapped dry before the addition of the tagged ligand (SARS-CoV-2 S1 or RBD) and test compounds diluted in binding buffer (100 mM HEPES, pH 7.2) to give a total volume of 100 μL/well. After 1 h incubation, three washes were conducted, and a further 1 h incubation with anti-His HRP conjugate diluted (1:2500) in SuperBlock (PBS) was used to detect the bound His-tagged ligand. Plates were washed four times before the addition of 100 μL/well of HRP substrate TMB (3,3′,5,5′-tetramethylbenzidine) and kept in the dark for up to 15 min. The reaction was stopped using 20 μL of 1M H_2_SO_4_, and the absorbance value was read at 450 nm. The plated concentrations of ACE2 receptor were 1.0 μg/mL for SARS-CoV-2 RBD and 2.0 μg/mL for SARS-CoV-2 S1. The concentrations of the ligand used in the inhibitory assays were 0.5 μg/mL for RBD and 1.0 μg/mL for S1. These values were selected following preliminary testing to optimize response (i.e., to produce a high-enough signal at conditions close to half-maximal response, EC_50_). Binding assessments for CD40–CD40L and TNF-R1–TNF-α were performed as previously described (Bojadzic et al., 2018). Stock solutions of compounds at 10 mM in DMSO were used.

### Statistics and Data Fitting

All binding inhibition and cell assays were tested in at least duplicate per plates, and assays were performed as at least two independent experiments. As before (Ganesan et al., 2011; Song et al., 2014; Chen et al., 2017), binding data were converted to percent inhibition and fitted with standard log inhibitor vs. normalized response models (Buchwald, 2020) using nonlinear regression in GraphPad Prism (GraphPad, La Jolla, CA, USA) to establish half-maximal (median) effective or inhibitory concentrations (EC_50_, IC_50_).

## RESULTS

As part of our work to identify SMIs for co-signaling PPIs that are essential for the activation and control of immune cells, we discovered that the chemical space of organic dyes, which is particularly rich in strong protein binders, can offer a useful starting point. Accordingly, it seemed logical to explore it for possible inhibitors of the SARS-CoV-2 S protein – ACE2 PPI that is an essential first step for the viral entry of this novel, highly infectious coronavirus. As a first step, we explored the feasibility of setting up a screening assays using a cell-free ELISA-type 96-well format similar to those used in our previous works with Fc-conjugated receptors coated on the plate and FLAG- or His-tagged ligands in the solution (Margolles-Clark et al., 2009b; Ganesan et al., 2011; Song et al., 2014; Chen et al., 2017). To establish assay conditions, we first performed concentration-response assessments using such a format with ACE2-Fc and SARS-CoV-2 S1 or RBD with His tag, and they indicated that both bindings follow classic sigmoid patterns with a slightly stronger binding for RBD than S1 (Figure 1). Fitting of data gave median effective concentrations (EC_50_s) and hence binding affinity constant (*K*_d_) estimates of 5 and 13 nM, respectively (127 and 1008 ng/mL) – in good agreement with the specifications of the manufacturer and published values that are also in the low nanomolar range (4–90 nM), typically based on surface plasmon resonance (SPR) studies (Sivaraman et al., 2020).

**Figure 1.**
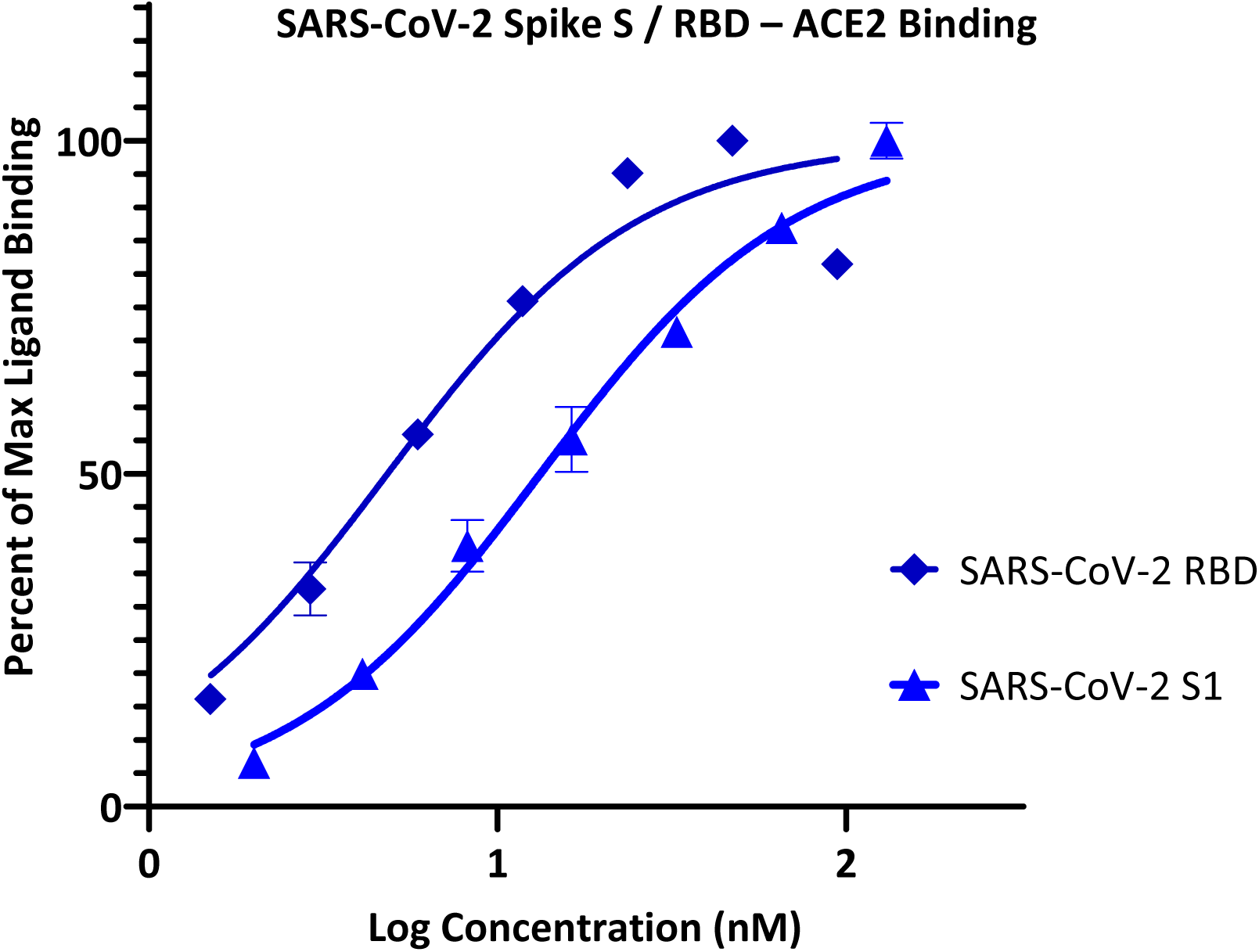
Concentration-response curves for binding of SARS-CoV-2 spike protein S1 and RBD to ACE2 in our ELISA-based assay format. Data obtained with Fc-conjugated ACE2 coated on the plate and His-tagged S1 or RBD added in increasing amounts as shown with the amount bound detected using an anti-His–HRP conjugate (mean ± SD for two experiments in duplicates).

Accordingly, we can use this format for inhibitory screening, and we decided to use hACE2 with SARS-CoV-2 RBD-His, as it showed stronger binding. In fact, this assay setup is very similar to one recently shown to work as a specific and sensitive SARS-CoV-2 surrogate virus neutralization test based on antibody-mediated blockage of this same PPI (CoV-S–ACE2) (Tan et al., 2020). With this setup in our hands, we performed a preliminary screening of representative organic dyes from our in-house library plus a few compounds that are or have been considered of possible interest in inhibiting SAR-CoV-2 by different mechanisms of action, e.g., chloroquine, clemastine, and suramin (Colson et al., 2020; da Silva et al., 2020; Gordon et al., 2020; McKee et al., 2020; Xiu et al., 2020). Screening at 5 μM indicated that most have no activity and, hence, are unlikely to interfere with the S-protein – ACE2 binding needed for viral attachment. Nevertheless, some showed activity; those of selected compounds of interest are shown in Figure 2 together with corresponding chemical structures. Erythrosine B (ErB, FD&C red #3), an FDA approved food colorant, was included as a possible positive control since we have shown it previously to be a promiscuous PPI inhibitor together with other xanthene dyes (Ganesan et al., 2011), and it indeed showed strong inhibition here. Of particular interest, methylene blue (MeBlu), which has a long history of diverse medical applications (Clifton and Leikin, 2003; Schirmer et al., 2011; Bistas and Sanghavi, 2020), also showed promising inhibitory activity.

**Figure 2.**
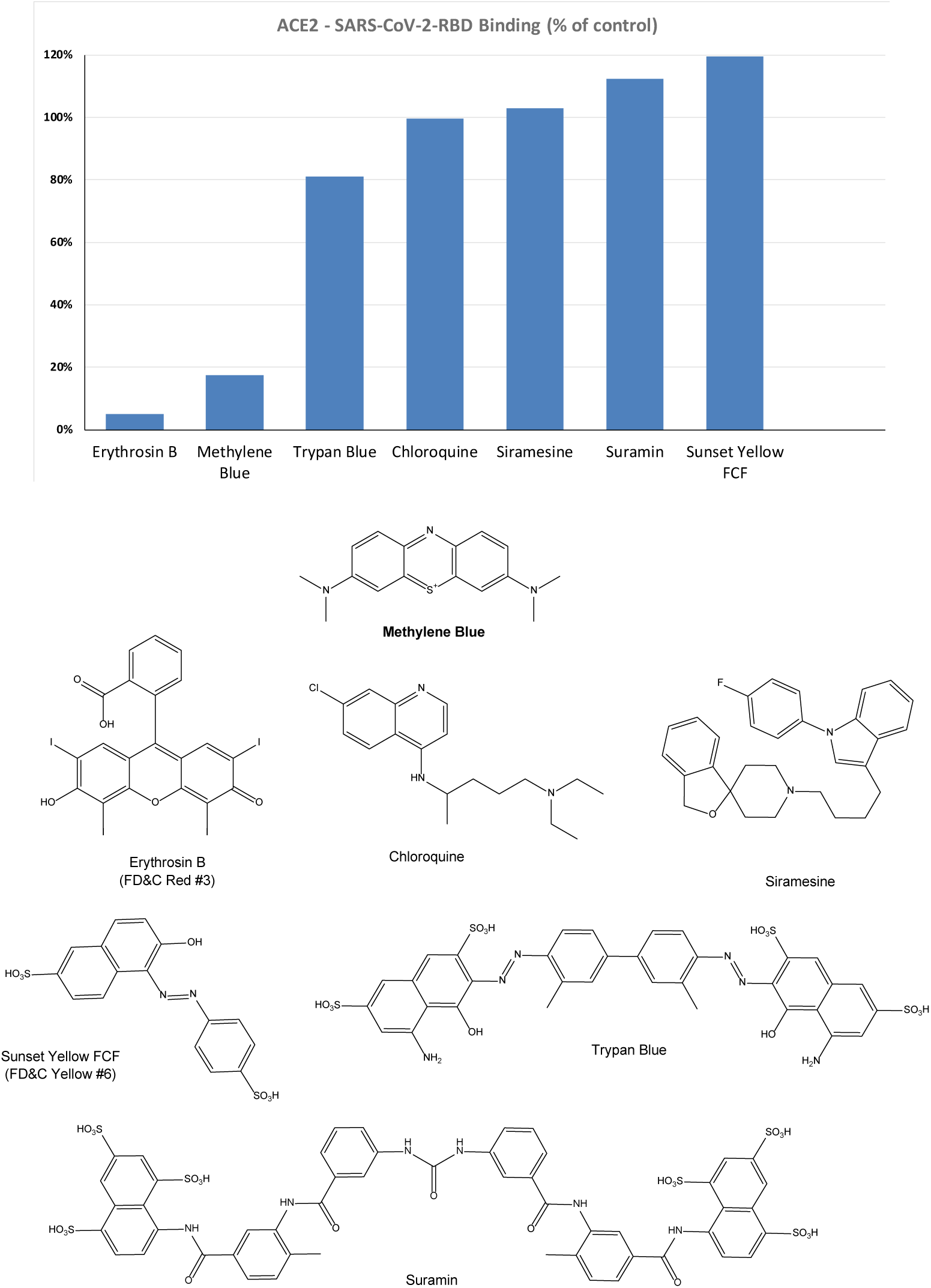
Inhibitory effect of selected compounds on SARS-CoV-2 RBD binding to hACE2 in our screening assay. Percent inhibition values obtained at 5 μM concentration shown normalized to control (100%). Erythrosine B, a known promiscuous SMI of PPIs (Ganesan et al., 2011) and sunset yellow FCF (FD&C yellow no. 6), a food colorant likely to be inactive, were included as positive and negative controls, respectively. Chemical structures are shown for comparison purposes.

Therefore, to confirm its activity, we performed detailed concentration-response assessments as recommended per experimental guidelines in pharmacology and experimental biology (Curtis et al., 2018; Michel et al., 2020). As shown in Figure 3, this confirmed that MeBlu indeed inhibits concentration-dependent manner an estimated IC_50_ of 3 μM, whereas chloroquine and suramin show no inhibitory activity in this assay. Chloroquine, an anti-parasitic and immunosuppressive drug primarily used to prevent and treat malaria, was included as it has potential antiviral activity against SARS-CoV-2 (subject to controversies) (Colson et al., 2020). Suramin, an antiparasitic drug approved for the prophylactic treatment of African sleeping sickness (trypanosomiasis) and river blindness (onchocerciasis), was incorporated because it was claimed to inhibit SARS-CoV-2 infection in cell culture most likely by preventing binding or entry of the virus (da Silva et al., 2020) (as well as because we found it earlier to inhibit the CD40–CD40L PPI (Margolles-Clark et al., 2009a)). ErB also inhibited with an IC_50_ of 0.4 μM, which is consistent with our previous observation of promiscuous PPI inhibition by this compound with a possibly slightly higher activity than found for other PPIs tested before (1–20 μM) (Ganesan et al., 2011). Sunset yellow FCF (FD&C yellow #6), an azo dye and an FDA approved food colorant included as a possible negative control, indeed showed no inhibitory activity. Since the IC_50_ obtained for MeBlu here (3 μM) is within the range of its circulating levels following normal clinical dosage (e.g., peak blood concentration of 19 μM after 500 mg p.o. with an elimination half-life of ∼14 h (Walter-Sack et al., 2009) or trough levels of 6–7 μM in healthy human volunteers following oral doses of 69 mg, t.i.d. = 207 mg/day (Baddeley et al., 2015)), this inhibitory effect on viral attachment can contribute to the possible antiviral activity of MeBlu against SARS-CoV-2 and possibly other ACE2-binding CoVs such as SARS-CoV and the α-coronavirus HCoV NL63.

**Figure 3.**
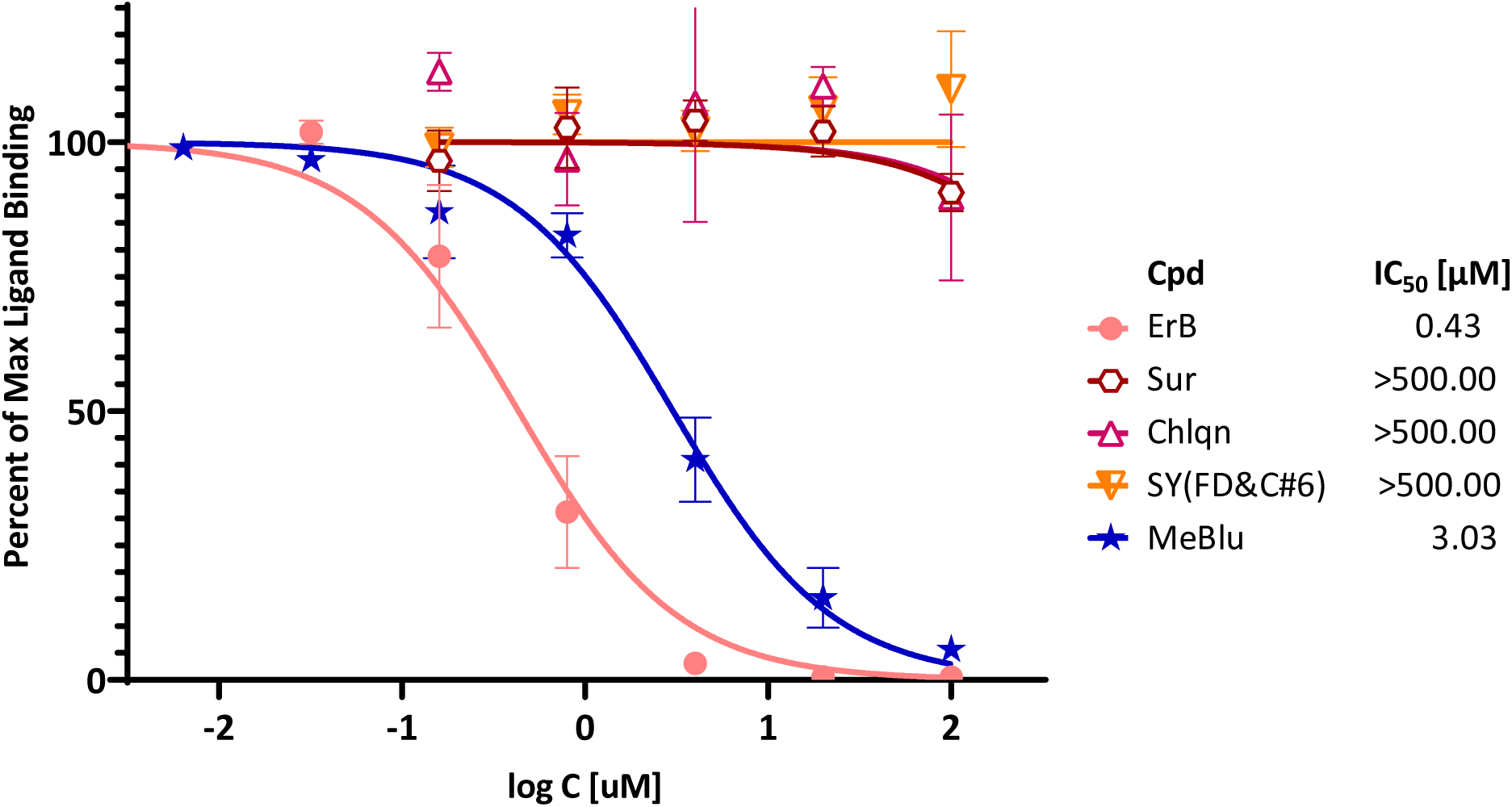
Concentration-dependent inhibition of SARS-CoV-2 RBD binding to ACE2 by selected compounds. Concentration-response curves obtained in ELISA-type assay with Fc-conjugated ACE2 coated on the plate (1 μg/mL) and His-tagged RBD (0.5 μg/mL) added and amount bound in the presence of increasing concentrations of test compounds detected. As before, erythrosine B (ErB) and sunset yellow FCF (SY(FD&C#6)) were included as positive and negative controls, respectively. Data (mean ± SD for two experiments in duplicates) were normalized and fitted with standard inhibition curves; obtained IC_50_ values are shown at right.

## DISCUSSION

Results here confirm again the usefulness of our strategy to rely on the chemical space of organic dyes, known to contain strong protein binders, as a starting platform to identify SMI scaffolds for PPI inhibition. Using this strategy, we have achieved considerable progress in targeting co-signaling interactions as we have identified the first SMIs for CD40–CD40L (Margolles-Clark et al., 2009b) and OX40– OX40L PPIs (Song et al., 2014) as well as the first promiscuous SMIs of PPIs (Ganesan et al., 2011). Organic dyes contain privileged structures for protein binding (Che et al., 2006; Fletcher and Hamilton, 2006; Hershberger et al., 2007), and, contrary to usual drug-like libraries, whose chemical space does not correspond well with that of promising PPI inhibitors (Neugebauer et al., 2007; Reynès et al., 2010; Sperandio et al., 2010), they are a good starting point to identify SMIs of PPIs. Most dyes, however, are unsuitable for therapeutic development because of their strong color and, in the case of azo dyes, their quick metabolic degradation (Levine, 1991; Feng et al., 2012); hence further medicinal chemistry is needed to optimize their clinical potential (Chen et al., 2017).

More importantly, our results indicate that MeBlu, an organic dye in clinical use for some therapeutic applications in the developed world (Clifton and Leikin, 2003; Schirmer et al., 2011; Bistas and Sanghavi, 2020) and with additional potential for certain developing world applications such as malaria (Dicko et al., 2018), can inhibit the viral attachment and entry of SARS-CoV-2 by blocking the PPI of its spike protein with ACE2 on the host cell. MeBlu is a tricyclic phenothiazine dye approved by the FDA for clinical use for the treatment of methemoglobinemia, and it is also used for other applications such as prevention of urinary tract infections in elderly patients; ifosfamid-induced neurotoxicity in cancer patients; vasoplegic syndrome, a type of distributive shock that occurs during coronary procedures; and intraoperative visualization of nerves, nerve tissues, and endocrine glands (Schirmer et al., 2011; Bistas and Sanghavi, 2020). MeBlu is included in the WHO List of Essential Medicines and was, in fact, the very first fully synthetic drug used in medicine, as it was used to treat malaria since 1891 (Schirmer et al., 2011). This utilization spanned through WW2 until it was replaced by chloroquine; although, due to the blue urine it could cause, MeBlu was not well liked among the soldiers (“Even at the loo we see, we pee, navy blue”) (Schirmer et al., 2011). It also served as the lead compound for the development of chlorpromazine and tricyclic antidepressants (Schirmer et al., 2011). Moreover, there is resurgent interest in its antimalarial application (Dicko et al., 2018), and it has potential for the treatment of neurodegenerative disorders such as Alzheimer’s disease (AD), due to its putative inhibitory action on the aggregation of tau protein (Schirmer et al., 2011). Notably, MeBlu was also part of the first method developed for pathogen inactivation in plasma, where it has been used since 1991 to inactivate viruses in combination with light (Lozano et al., 2013). MeBlu intercalates within nucleic acid strands, and application of light causes its excitation generating highly reactive singlet oxygen that oxidizes guanosine and breaks nucleic strands (Lozano et al., 2013). Hence, in the presence of light, MeBlu has broad-spectrum virucidal activity and is used to inactivate viruses in blood products prior to transfusions.

Notably, there is also recent evidence of possible *in vitro* antiviral activity for MeBlu even in the absence of UV-induced activation, as MeBlu showed virucidal activity at low micromolar concentrations when incubated with SARS-CoV-2 for 20 h in the dark (Cagno et al., 2020). Its ability to inhibit the SARS-CoV-2-S – ACE2 PPI could be a mechanism of action contributing to such activity. If this PPI inhibitory activity of MeBlu is retained at similar levels *in vivo* as found here in a cell-free assay (IC_50_ = 3 μM), it is within a range that can be obtained in blood following typical doses (e.g., 200 mg/day) as indicated by pharmacokinetic studies in humans. For example, in one study, peak blood concentration of MeBlu was 19 μM after 500 mg p.o., and the elimination half-life was also more than adequate being around 14 h (Walter-Sack et al., 2009). In another study, trough levels of 6–7 μM were obtained following total daily oral doses of 207 mg/day (administered as 69 mg, p.o., t.i.d.) (Baddeley et al., 2015). Hence, oral administration could provide adequate concentrations (e.g., > 7 μM) and inhaled applications, which have been explored in less developed countries for some respiratory treatments (Golwalkar, 2020), could be even more advantageous. MeBlu is generally safe, but it shows dose-dependent toxicity with nausea, vomiting, hemolysis, and other undesired side effects starting to occur at doses >7 mg/kg (i.e., >500 mg) (Clifton and Leikin, 2003; Bistas and Sanghavi, 2020). It also is contraindicated in certain populations, e.g., in those taking serotonin reuptake inhibitors and in persons with hereditary glucose-6-phosphate dehydrogenase deficiency (G6PD deficiency) (Schirmer et al., 2011; Bistas and Sanghavi, 2020).

It has to be noted, however, that MeBlu also inhibited the CD40–CD40L and TNF-R1–TNFα PPIs in our assays with low- to mid-micromolar potency (data not shown); hence, it is possible that MeBlu is a somewhat promiscuous PPI inhibitor limiting its usefulness. Its three-ring phenothiazine framework resembles somewhat the three-ring xanthene framework of erythrosine B (Figure 2), which we have shown before to act as promiscuous PPI inhibitor together with some other structural analog xanthene dyes such as rose Bengal and phloxine (Ganesan et al., 2011). MeBlu certainly shows polypharmacology and acts on a multitude of targets (Schirmer et al., 2011); many of these however can have further beneficial effects in COVID-19 patients. Its main mechanism of action is reducing the oxidized ferric form of hemoglobin (Fe^3+^) when in a state of methemoglobinemia, which binds oxygen irreversibly, to the ferrous (Fe^2+^) form (Bistas and Sanghavi, 2020). This increases the oxygen-binding capacity of hemoglobin and, thus, oxygen delivery to tissues – an important benefit for COVID-19 patients. COVID-19 patients often exhibiting low oxygen levels, typically incompatible with life without dyspnea – a phenomenon termed silent hypoxemia (or happy hypoxia in public media) (Tobin et al., 2020). Possibly relevant to this, MeBlu was found to improve hypoxemia and hyperdynamic circulation in patients with liver cirrhosis and severe hepatopulmonary syndrome (Schenk et al., 2000). MeBlu is being used for the treatment of pneumonia and other respiratory ailments in less developed countries with some success (Golwalkar, 2020).

Further, MeBlu was recently shown to block the PD-1–SHP2 PPI, which is downstream from the PD-1–PD-L1 co-signaling PPI, with low micromolar potency and effectively enough to counteract the suppressive activity of PD-1 on cytotoxic T lymphocytes and restore their cytotoxicity, activation, proliferation, and cytokine-secreting activity (Fan et al., 2020). This mechanism of action targeting this co-signaling pathway (PD-1) could contribute to restoring T cell homeostasis and function from exhausted state (Barber et al., 2006; Vardhana and Wolchok, 2020), which is of interest to improve viral clearance and rein-in the inflammatory immune response and the associated cytokine storm during anti-viral responses such as those causing the high mortality of COVID-19 patients (Liu et al., 2020; Ye et al., 2020).

As far as clinical applications, one promising indication comes from a report of a cohort of 2500 French patients treated with MeBlu as part of their cancer care none of whom developed influenza like illness during the COVID-19 epidemics (Henry et al., 2020). MeBlu has also been explored in one Phase 1 clinical trial (NCT04370288) for treatment of critically ill COVID-19 patients in Iran as part of a three-drug last therapeutic option add-on cocktail (MeBlu 1 mg/kg, vitamin C 1500 mg/kg, and N-acetyl cysteine 2000 mg/kg) based on the hypothesis that this combination could rebalance NO, methemoglobin, and oxidative stress. Four of the five patients responded well to treatment (Alamdari et al., 2020).

In conclusion, screening of our organic dye-based library identified MeBlu as a low-micromolar inhibitor of the interaction between SARS-CoV-2 spike protein and its cognate receptor ACE2, a PPI that is the first critical step initiating the viral entry of this coronavirus. While MeBlu shows strong polypharmacology and might be a somewhat promiscuous PPI inhibitor, its ability to inhibit this PPI could contribute to the antiviral activity of MeBlu against SARS-CoV-2 even in the absence of light making this inexpensive and widely available drug potentially useful in the prevention and treatment of COVID-19 as an oral or inhaled medication.

## ABBREVIATIONS

ACE2: angiotensin converting enzyme 2;
CoV: coronavirus;
MeBlu: methylene blue;
PPI: protein-protein interaction;
SARS: severe acute respiratory syndrome;
SMI: small-molecule inhibitor.

## ACKNOWLEDGEMENTS

Financial support by the Diabetes Research Institute Foundation (www.diabetesresearch.org) is gratefully acknowledged.

## AUTHOR CONTRIBUTIONS

DB performed most of the experiments, OA contributed to some. PB originated and designed the project, provided study guidance, and wrote the draft manuscript. All authors contributed to writing and read the final manuscript.

